# SnapG: An Automated Tool for Myelin g-ratio Measurement with Batch Processing Capability

**DOI:** 10.64898/2026.01.14.699516

**Authors:** Yoko Bekku, Stephen Cai, Thomas Xin, Joseph Sall, Frances Kestel, Alice F Liang, James L. Salzer

## Abstract

The g-ratio, calculated as the axon diameter divided by the total myelinated fiber diameter, is widely used to assess the degree of myelination in the central and peripheral nervous systems. Changes in g-ratios accompany demyelinating, hypomyelinating, and remyelinating conditions, and can also result from activity-dependent, adaptive myelination. Although manual tracing from electron micrographs has been the gold-standard to measure g-ratios, this process is laborious and limits throughput. To address this challenge, we developed SnapG, an automated and user-friendly software tool to streamline g-ratio analysis from electron micrographs. SnapG implements an end-to-end pipeline for axon and myelin segmentation on EM images. The tool integrates robust segmentation and post-processing steps to extract axon and fiber diameters and generates both per-image and cohort-level summary statistics. SnapG produces g-ratio distributions that agree well with expert manual annotations and established tools across imaging modalities. It reduces user intervention, supports large image and batch processing, and achieves faster processing times while delivering accurate, reproducible measurements. SnapG’s automated reporting thereby facilitates high-throughput analyses and improves consistency across experiments. SnapG can accelerate studies of myelin biology and provides a practical alternative to existing tools.

## Introduction

Myelin, the multilamellar glial sheath that surrounds axons, emerged during evolution some 425 million years ago in the earliest hinge-jaw fishes (Zalc, 2016). The appearance of myelin made the development of the large, complex nervous systems of vertebrates possible. In addition to its key role in enabling saltatory conduction, myelin is now appreciated to have critical roles in the metabolic support of axons and in optimizing circuit function via activity-dependent, adaptive changes (Monje, 2018; Simons et al., 2024).

Since Remak’s initial description of myelin (Remak, 1839), it has been the subject of intense efforts to characterize its function and assess its structure. By using birefringence of images obtained through optical microscopy, Schmitt and Bear were the first to quantify the thickness of myelin (Bear and Schmitt, 1937). Their studies defined the g-ratio (initially termed g) as the ratio of the axon diameter to the outer diameter of the myelin sheath (Gow, 2025). The g-ratio has since become the standard metric of myelin thickness with values of ∼ 0.67 in the peripheral nervous system (PNS) and ∼ 0.77 in the central nervous system (CNS). These values are generally conserved across species and for different PNS nerves or different white matter tracts in the CNS. The measured values in the PNS agree well with an optimal g-ratio of ∼0.6 that supports maximal conduction speed as theoretically derived by Rushton (Rushton, 1951). The higher g-ratios in the CNS may reflect additional constraints, including intracranial volume conservation (Chomiak and Hu, 2009). Alterations in g-ratios are common in neurological disorders accompanied by hypomyelination, demyelination +/− remyelination such as Charcot-Marie-Tooth disease and multiple sclerosis underscoring the importance of accurate g-ratio measurements. In addition, g-ratios can also be modulated normally, for example reflecting increases in myelin thickness during activity-dependent plasticity (Fields, 2015; Mount and Monje, 2017).

Despite its broad utility as a measure of myelin thickness, g-ratio quantification remains a time-consuming process that traditionally requires extensive manual tracing of axon and myelin profiles from electron micrographs. To address the challenges of manual g-ratio quantification, several automated tools have been developed in recent years to improve efficiency, accuracy, and data consistency; examples include MyelTracer, AxonSeg, and AxonDeepSeg (Kaiser et al., 2021; Zaimi et al., 2016; Zaimi et al., 2018). These tools are compatible with either electron micrographs or optical microscopic images, depending on the software. They operate in a semi-automated manner. AxonDeepSeg, a deep learning-based axon and myelin segmentation tool, produces highly accurate results across various species and works with both SEM and TEM images (Zaimi et al., 2018). However, it typically requires powerful hardware and longer processing times to achieve a high level of accuracy. MyelTracer, on the other hand, relies on traditional image processing techniques and lower-resolution images, but requires supervision by the user during segmentation (Kaiser et al., 2021). This limits the range of usable images and makes large-scale batch processing difficult.

To further enhance efficiency and reduce user burden, there is a remaining need for fully automated software that eliminates pre-processing requirements, supports unrestricted image sizes, and offers a user-friendly interface. Here, we describe SnapG, an automated tool we have developed for myelin g-ratio analysis from electron micrographs

## Materials and Methods

### Mice

C57BL/6 mouse line was used in this study. Mice were sacrificed at 3 months of age. All experiments with mice were performed in compliance with the relevant policies and institutional guidelines issued by the New York University School of Medicine Institutional Animal Care and Use Committee.

### Electron microscopy

Mice were transcardially perfused with 4% PFA, 2.5% glutaraldehyde and 0.1 M sucrose in 0.1 M cacodylate buffer (CB), pH 7.4. After an additional 2 hrs, sciatic nerves, optic nerves, and whole brains were dissected and immersed in the same fixative solution overnight. The genu of corpus callosum was dissected by a brain slicer. Fixed tissues were washed three times in 0.1% CB for 15 minutes each, then post-fixed in 1% osmium tetroxide in CB for 2 hours at room temperature. The tissues were dehydrated in a series of ethanol solutions including 30, 50, 70, 85, 95, 100% and 100% for 15 min in each solution at room temperature, then immersed in propylene oxide (PO) (Electron Microscopy Sciences #20412), 1:1 solution mix of PO: Epon without BDMA (46% EMbed 812 (Electron Microscopy Sciences #14900) + 36% DDSA (Electron Microscopy Sciences #13710) + 18% NMA (Electron Microscopy Sciences #19000)) for 1 hr, 1:2 of solution mix of PO: Epon without BDMA overnight, and embedded in Epon (44% EMbed 812 + 35% DDSA + 18% NMA + 3% BDMA (Electron Microscopy Sciences #14900). Samples were sectioned at 70 nm thickness using Leica UC6 ultramicrotome (Germany) and stained with uranyl acetate and lead citrate as standard manner. Electron micrographs of random fields were captured at the NYU Langone Microscopy Laboratory on a JEOL JEM-1400 Flash TEM (Japan) and photographed with a Gatan Rio16 camera (Gatan Inc.), or a Talos L120 C TEM (Thermo Fisher Scientific) coupled with a Gatan OneView camera. All chemicals and EM grids are purchased from Electron Microscopy Sciences, Hatfield, PA.

### Software development

SnapG relies on the Python NumPy and OpenCV libraries and consists of two main steps: preprocessing and information extraction. In the preprocessing step, SnapG reads the input image in grayscale, where each pixel’s brightness is represented by an integer value from 0 (black) to 255 (white). The image can then be resized to dimensions smaller than the original to increase processing speed and reduce noise; users may choose the appropriate size based on their computer’s performance. To further suppress noise, SnapG applies an average filter using a circular kernel, defined by a user-specified integer radius *r* and generated with numpy.ogrid:

import numpy as np
import cv2
y, x = np.*ogrid*[-r:r+1, -r:r+1]
kernel = x**2 + y**2 <= r**2
kernel /= np.*sum*(kernel)
img_gray= cv2.*filter2D* (src=img_gray, ddepth=-1, kernel=kernel)

The filtered image is then thresholded to a binary mask using a user-defined *threshold_val*, and small black features that individually take up less than 0.06% of the entire image (either image noise or small organelles) are filled in:

_, thresh = cv2.*threshold*(img_gray, threshold_val, 255, cv2.*THRESH_BINARY*)
inverted = cv2.*bitwise_not*(thresh)
contours, _ = cv2.*findContours*(inverted, cv2.*RETR_TREE*,
cv2.*CHAIN_APPROX_NONE*)
for c *in* contours:
area_proportion = cv2.*contourArea*(c) / total_image_area
if area_proportion < 0.0006:
cv2.*drawContours*(thresh, [c], −1, 255, cv2.*FILLED*)

A dilation operation with a user-defined integer kernel size *dilate_val* is applied, followed by a similar erosion with *erode_val*:

dilate_kernel = cv2.*getStructuringElement*(cv2.*MORPH_RECT*, (dilate_val,dilate_val))
dilated = cv2.*morphologyEx*(thresh, cv2.*MORPH_DILATE*, dilate_kernel)
erode_kernel = cv2.*getStructuringElement*(cv2.*MORPH_RECT*, (erode_val,erode_val))
eroded = cv2.*morphologyEx*(dilated, cv2.*MORPH_ERODE*, erode_kernel)

Contours—outlines of contiguous groups of white pixels—are detecting using cv2.*findContours*. Invalid contours are excluded based on the following criteria: touching the image edges; bounding box area outside of a user-defined range [*min_area*, *max_area*] in pixels^2; or insufficient convexness or circularity, calculated as: convexness=convex_hull_area/contour_area circularity=4π⋅contour_area/(contour_perimeter)^2.

Contours that fail to meet the user-defined limits *min_convexness* and *min_circularity* are excluded. Remaining contours are assumed to represent axon inner edges, and the algorithm proceeds to the information extraction step. To approximate the myelin thickness for each contour, a mask of the contour is created and a distance transform is applied, mapping each pixel to its shortest Euclidean distance from the contour (Fig. S1A-C):

contour_mask = np.*zeros_like*(thresh)
cv2.*drawContours*(contour_mask, [current_contour], −1, 255, thickness=cv2.*FILLED*)
distance_transform = cv2.*distanceTransform*(255 - contour_mask,
distanceType=cv2.*DIST_L2*, maskSize=5)

An exclusion mask and an outline mask are created. The exclusion mask is the same as the preprocessed image, but it excludes the current contour. The outline mask contains the edges of the exclusion mask, derived using cv2.*Canny*:

exclusion_mask = (contour_mask == 0) & (eroded == 255)
outline_mask = cv2.*Canny*(exclusion_mask,0,0)

The outline mask is applied to the distance transform (Fig. S1D-F):

distances = distance_transform[outline_mask > 0]
distances = distances[distances > 0]

To extract myelin thickness, the *n* closest nonzero pixels to the contour are sampled, where *n* is twice the number of points in the current contour. From the resulting distribution of distances, the value at the *thickness_percentile* percentile is chosen as the representative myelin thickness in pixels, where *thickness_percentile* is a user-defined parameter (Fig. S1G):

= 2 * len(current_contour)
smallest_n = np.*partition*(distances, n – 1)[:n]
= np.*percentile*(smallest, thickness_percentile)

The axon’s inner radius is approximated as:

contour_perimeter = cv2.*arcLength*(inner_contour[0], closed=*True*)
radius=(contour_perimeter)/2π

The g ratio is then calculated:

g_ratio = radius / (radius + thickness)

Other measurements, such as inner and outer diameter, are derived from these approximations.

### Quantification of g-ratio in electron micrographs

For comparison between manual and SnapG-based g-ratio calculations, we used images originally acquired for a previous study from the lab (Bekku et al., 2024) together with the corresponding manually quantified g-ratio data. Briefly, manual g-ratio quantification was performed by calculating diameters from their respective perimeters as measured with the magnetic lasso tool of Photoshop (Adobe). For comparison between per-image and batch-level g-ratio calculation, images of the P90 carpus callosum were randomly acquired from a section on a grid, ensuring coverage across the entire region. For P90 sciatic nerves, 10 FOVs (1 FOV = 22.56 x 22.56 µm) per mouse were analyzed from 2 mice. For P90 optic nerves, 10 FOVs (1 FOV = 6.92 x 6.92 µm) per mouse were analyzed from 2 mice. For P90 corpus callosum, 26 FOVs (1 FOV = 6.78 x 6.78 µm) were analyzed from 1 mouse. Only axons with a circularity equal or greater than 0.6 were included in the analysis. SnapG can run on Windows 11 (Intel Core i5-14500, 32 GB RAM, Intel UHD Graphics 770; Intel Core i7-12700H, 16 GB RAM, Intel Iris Xe and NVIDIA GeForce RTX 3060 Laptop GPU; AMD Ryzen 7 8840HS, 16 GB LPDDR5 RAM, AMD Radeon Graphics), macOS devices including the MacBook Air (M1, 2020) and Mac mini (M2, 2023; macOS Sequoia), and Linux Mint v22.2 Cinnamon (AMD Ryzen 7 8840HS, 16 GB LPDDR5 RAM, AMD Radeon Graphics).

### Statistical analysis

Figure legends contain the number of samples and additional details of the data plotted. The results section also describes the number of axons, animals, or FOVs analyzed. All statistical significances were assessed by Student’s *t* test in Excel (Microsoft 365).

### Procedure

To begin, download the SnapG code, which is freely available online at https://github.com/KK201431873/SnapG. Open the terminal in the SnapG folder and run the software.

SnapG analysis consists of three steps:

1. Data Preparation — Parameter Tuning: Adjusting segmentation parameters and thickness percentiles as part of data preparation. This step is responsible for threshold adjustment and contour detection.
2. Reviewing Segmentation Files: Manually checking and correcting the segmentation results.
3. Data Generation: Performing quantification tasks, including g-ratio and myelin-thickness measurements.

Detailed procedures are also available on the SnapG website.

### Step 1. Data Preparation — Parameter Tuning

Before performing data analysis, users should define the thresholds that will be used for quantification.

1. Open image file(s) from the File menu (Fig. 1A).
2. Select the image(s) in the file dialog and click “Open.”
3. Tune the segmentation parameters in the “Segmentation Settings” panel.

a. Set the scale per pixel in “Dist. per pix.” You can choose either µm or nm.
b. Adjust the “Image Res. Divisor” value as needed. We usually use a divisor of 3 to downsample the images, but you may increase it up to 10.
c. Click “Show Threshold” to display the black-and-white thresholded image. Adjust the myelin-thickness percentile in “Thickness %tile.” We generally use 60% but you may need to determine the value best fits your samples. Keep this value consistent within the same dataset.
d. Adjust “Threshold,” “Radius,” “Dilate,” and “Erode” until the black-and-white image looks representative of your sample (Fig. 1B).
e. Click “Show Text” to display the tracing lines, axon IDs, and G-ratios on the original image. The corresponding data will appear in the output fields (Fig. 1C).
4. (Optional: Batch processing) Open the Batch Processing panel from the View menu, and click “Choose images” under “Data Preparation”.
5. Click “Add images” and select the image(s) for which you tuned the segmentation parameters.
6. Check the “Check selected” box and click “OK.”
7. Click “Choose destination path” and select the folder where the segmentation files will be saved.
8. Check “Use multiprocessing” to allow the computer to use the maximum number of logical processors.
9. Click “Start” in the “Processing Output” section.
10. Check the output folder. It will contain the “xxx.seg” file(s).

**Figure 1.**
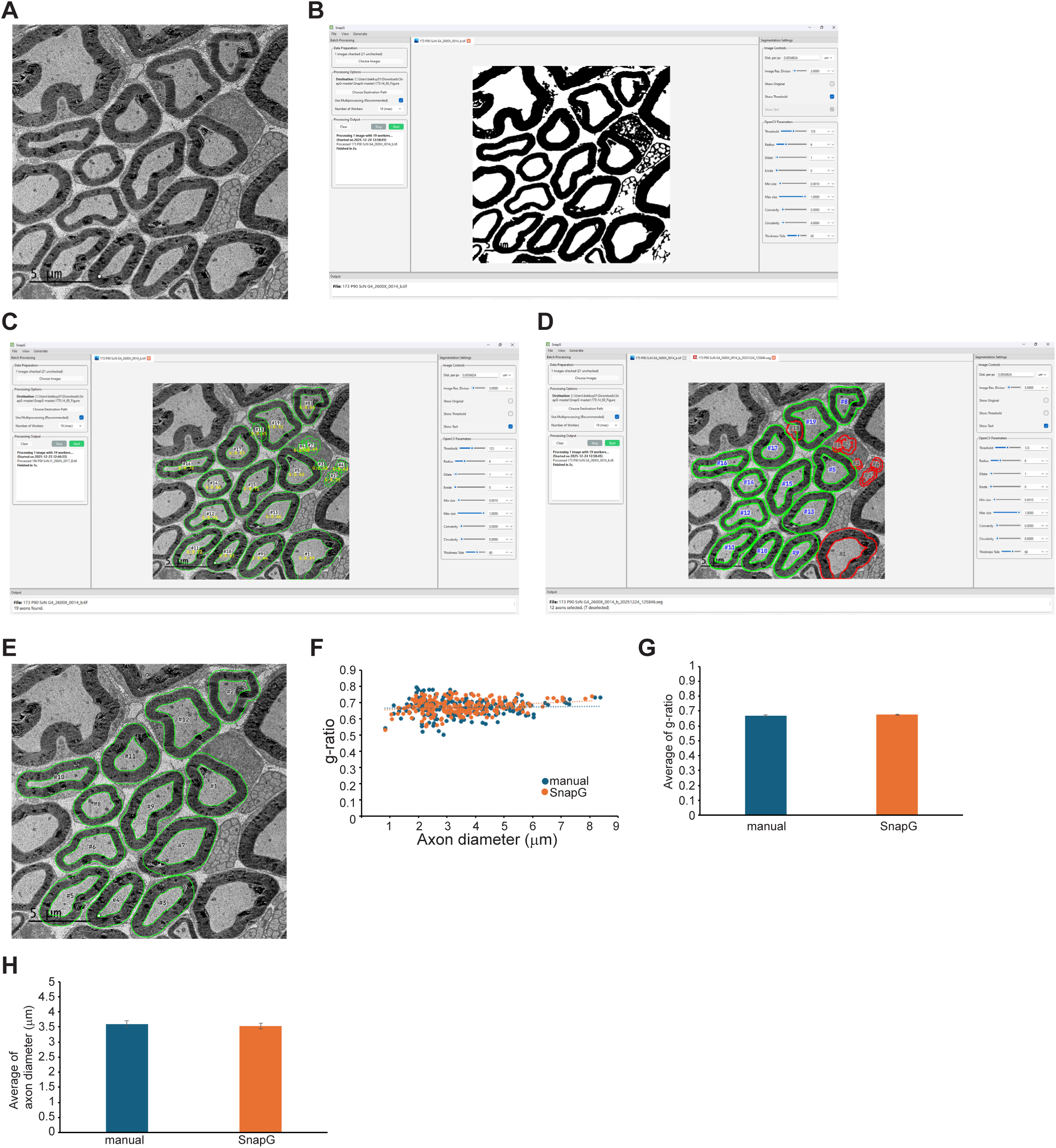
g-ratio calculated by SnapG vs. manual measurements of PNS myelinated axons. A. Electron micrographs of P90 sciatic nerves. Bar, 5 μm. B. The SnapG software interface showing the threshold view with adjusted parameters. C. The outer, inner contours and axon diameters are shown in the processed image. D. Processed image after axons have been deselected. Deselected axons are marked by red lines. E. The final output generated by the code. Axon numbers match those listed in the CSV. F. Comparison of g-ratio values obtained from manual (n = 201 axons) and SnapG (n = 201 axons) calculations show both methods yield similar results. G. Bar graph showing the average *g*-ratio calculated manually (0.669 ± 0.004; mean ± SEM) and by SnapG (0.675 ± 0.003). H. Bar graph showing the average axon diameter calculated manually (3.600 ± 0.101 μm; mean ± SEM) and by SnapG (3.529 ± 0.098).

### Step2. Reviewing Segmentation Files

While SnapG is designed to avoid selecting myelinated axons that are only partially included in the image, it may occasionally select them. Axons within Remak bundles may also be included, although this is rare. In this step, users can manually filter out any such axons as well as any traced regions that are not axons.

1. Click “Open → Segmentation file(s)” from the File menu and open the “xxx.seg” file(s).
2. Click the axon(s) you want to deselect. The tracing lines will turn red instead of green Fig. 1D).
3. (Optional: Batch processing) Repeat step 2 for all “xxx.seg” files.

### Step 3. Generation of data

In this step, the segmentation results are converted into a readable format.

1. Click “segmentation data” from the file menu.
2. Click “add files”, choose “xxx.seg” file(s), click “select all”, and check “check selected” box.
3. Click “generate”. SnapG will generate the data and open the folder once the data is generated.
4. The output CSV includes lists of axons data for each image with each axon ID corresponding to a number drawn in that image (Fig. 1E).

### Save and load setting

SnapG can save the current settings—including threshold values—and load them later.

Saving:

Click “Save → Current settings” from the File menu. The settings will be saved as a “xxxx.snpg” file.

Loading:

Click “Open → Settings file” from the File menu, select the “xxx.snpg” file, and open it.

## Results

### SnapG is an automated tool for myelin g-ratio analysis: setting threshold and percentiles

To streamline g-ratio analysis and improve data reliability, we developed SnapG, an open-source, automated, Python-based tool for g-ratio quantification that is compatible with macOS, Linux, and Windows operating systems. As SnapG facilitates batch-wise g-ratio computation, it significantly streamlines the overall analysis workflow. As a threshold-based tool, SnapG achieves high detection rates of myelinated axons and substantially reduces processing time. SnapG thus provides a practical solution for large-scale g-ratio quantification in myelin research.

SnapG is composed of three steps as described in the procedure section. The identification of inner and outer contours is driven by the thresholds defined in the first step. SnapG displays both the threshold image and the detected contours overlaid on the original image. In the processed image, the user can define the pixel-to-nanometer conversion, enabling accurate measurement of inner contour diameters. Additionally, overlaying the contours on the original image allows users to adjust the threshold with precision (Fig. 1A -E).

To quantify myelin thickness, SnapG traces the inner contour and, at each point along it, locates the nearest white pixel outside the contour. Users specify a percentile value for averaging myelin thicknesses, and the distances at that percentile are used as the representative thickness. Percentiles are chosen to reduce the impact of outliers and to provide a more robust estimate of typical myelin thickness. In densely packed regions of the CNS, this procedure may inadvertently capture the inner contours of adjacent myelin sheaths. Consequently, the resulting outer contours overlaid on the original image can appear larger than the true outer boundaries. Selecting a smaller percentile (∼30%) helps to limit these enlarged outer contours, facilitating separation of two closely apposed myelinated axons whose boundaries may otherwise be obscured due to limitations in threshold-based contour detection. In contrast, in images where axons are well separated, such as in the PNS, larger percentiles (60 – 70%) may yield slightly more precise myelin thickness measurements.

The CSV output of SnapG provides detailed information, including the total number of axons detected, individual axon IDs, g-ratio, circularity, inner and outer contour diameters (μm or nm), and myelin thickness (μm or nm). Circularity was calculated using formula 4pi (area/perimeter^2^); axons with circularity values below 0.6 are typically excluded from g-ratio analyses in our lab. However, these axons are retained in the CSV output and are included in the total count of detected axons. This allows for meaningful comparisons of myelinated axon counts per FOV across normal and abnormal conditions.

### SnapG reproduces manual g-ratio quantifications in PNS

To assess the accuracy of SnapG, we calculated g-ratios with SnapG using electron micrographs of P90 sciatic nerve samples we had previously used in manually measured g-ratios (Bekku et al., 2024), selecting the same 201 myelinated axons for comparison. In the PNS, inner contours were interpreted as corresponding effectively to axon boundaries. First, we compared the 30th to 70th percentiles under the same parameters, including the threshold, to determine which percentile yields a g-ratio most similar to that determined by manual tracing (0.669 ± 0.0044; mean ± SEM). G-ratios calculated using the 30% or 40% were similar but slightly higher (30%: 0.698 ± 0.003, *p* < 0.001; 40%: 0.691 ± 0.003, *p* < 0.001) whereas those calculated using the 60% (0.675 ± 0.003, *p* = 0.25) or 70% (0.667 ± 0.003, *p* = 0.64) were equivalent with no significant differences (Fig. S1H). A scatter plot of g-ratio versus axon diameter is shown in Figure 1F. There is no significant difference between manual and SnapG measurements at 60% (Fig. 1G). The average axon diameter also showed no significant difference (manual: 3.596 ± 0.1011 μm, SnapG: 3.529 ± 0.098, *p* = 0.63) (Fig. 1H). These results indicate that SnapG is useful for high-throughput quantification in the PNS.

### SnapG recapitulates manual g-ratio measurements in the CNS

To evaluate the accuracy of SnapG in calculating g-ratios in the CNS, we re-analyzed P90 optic nerve samples from our previous study (Bekku et al., 2024), comparing manually obtained measurements with those generated by SnapG. In the CNS, a periaxonal space containing adaxonal myelin is sometimes observed between the axon and its surrounding myelin (Fig. 2A - C). In automated g-ratio quantification using SnapG, axons with a periaxonal space must either be excluded by the user or manually measured using custom contour definitions if inclusion is desired. Most axons were identified based on their inner contours, interpreted as axonal boundaries, while outer contours were used to define the outermost layer of the myelin sheath. In our previous study, axon diameters were measured manually; therefore, to ensure consistency between data sets, axons with periaxonal space were manually measured in this study as well (Fig. 2C). The outer myelin contours were determined using the 30% of the myelin-thickness measurements. Manually measured axons represented 24.61 ± 2.16 % of all myelinated axons (18 FOVs; mean ± SEM). The average g-ratio obtained from manual and SnapG measurements showed no significant difference (manual: 0.748 ± 0.003; mean ± SEM; SnapG: 0.757 ± 0.004, *p* = 0.132). Similarly, the average axon diameter did not differ between methods (manual: 0.835 ± 0.020 μm, SnapG: 0.839 ± 0.018, *p* = 1.96). The number of myelinated axons detected (circularity > 0.6) was largely consistent between manual and SnapG qualifications, with SnapG mode identifying 125.96 ± 7.13 % of the axons detected manually (manual: 245 axons; SnapG: 273 axons). These results indicate that SnapG provides reliable large-scale quantification of CNS myelinated axons.

**Figure 2.**
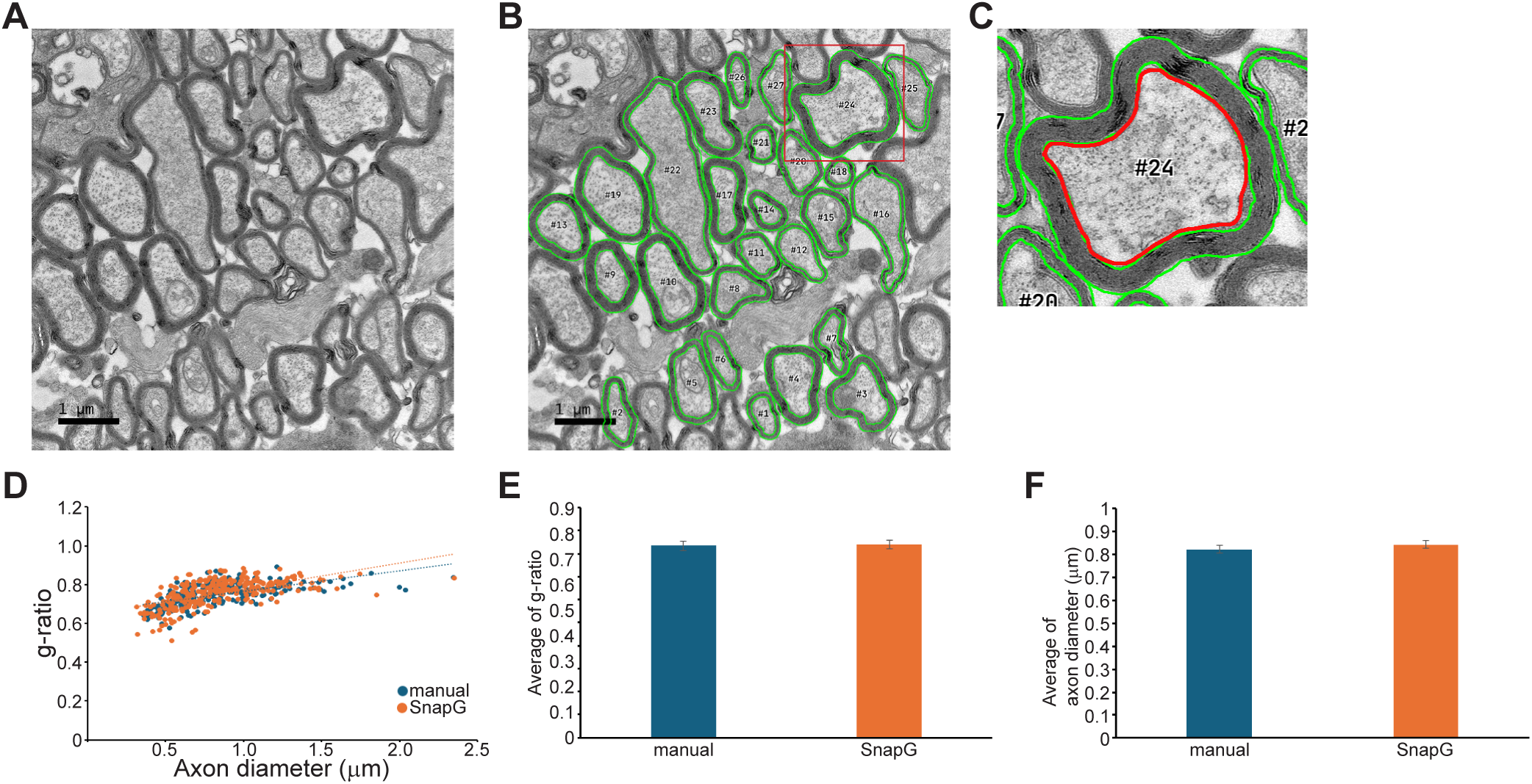
g-ratio calculated by SnapG vs. manual measurements of CNS myelinated axons. A. Electron micrographs of P90 optic nerves. Bar, 1 μm. B. The final output generated by the code. Axon numbers match those listed in the CSV. C. Higher magnification of the red rectangle shown in B. The red line in C indicates a manually traced axon. Notably, the traced line was drawn to exclude the periaxonal space. D. Comparison of g-ratio values obtained from manual (n = 245 axons) and SnapG (n = 273 axons) calculations shows both methods yield similar results. E. Bar graph showing the average g-ratio calculated manually (0.748 ± 0.004) and by SnapG (0.757 ± 0.004). F. Bar graph showing the average axon diameter calculated manually (0.835 ± 0.020 μm) and by SnapG (0.839 ± 0.018 μm).

### SnapG enables time-efficient g-ratio quantification via batch processing

Quantification of g-ratios remains a labor-intensive process, primarily due to the need for manual tracing across numerous randomly selected images per animal. This challenge is exacerbated in regions with heterogeneous distributions of myelinated axons, such as the corpus callosum, which necessitate increased sampling to obtain representative measurements. SnapG streamlines this process by enabling either individual or batch-wise image g-ratio calculations. In the individual image mode, each image is thresholded separately. In the batch quantification mode, a series of images captured with consistent brightness and contrast setting are thresholded and quantified together providing additional time savings. For example, once batch quantification is initiated at the data-generation step, SnapG completes the output generation typically within one minute, placing all output files into a single folder. This folder contains the multiple TIFF images with overlaid contours of selected axons. It also contains a single CSV file summarizing the numbers of myelinated axons detected, individual axon IDs, g-ratio, circularity, inner and outer contour diameters (nm or μm), and myelin thickness (nm or μm). This streamlined format facilitates downstream analysis and enables efficient comparison across datasets or experimental conditions.

We assessed the consistency of SnapG’s output between individual and batch quantification modes in the corpus callosum of P90 mice. To compare quantification of individual images vs. the batch-mode processing mode, we excluded the periaxonal space to consistently define the inner contours as axonal boundaries. We found the average g-ratios were quite similar between individual and one-batch measurements, although the slight differences were nevertheless significant (individual: 0.819 ± 0.001, mean ± SEM; batch: 0.826 ± 0.001; *p* < 0.001). The average axon diameter showed no significant differences (individual: 0.729 ± 0.007 μm; batch: 0.738 ± 0.008 μm; *p* = 0.43).

Thus, batch-processing represents a significant reduction in the time required to analyze g-ratio across multiple animals albeit with a slight compromise in the accuracy of g-ratio measurements. For the most accurate measurements, individual image analysis is recommended; for fastest analysis, batch-processing is recommended. Finally, we used a hybrid, multi-batch strategy of dividing the image series (26 images) into seven sub-groups, which were then batch-processed separately. In this analysis, there were no significant differences between individual and batch measurements in either the average g-ratio or the average axon diameter (Fig. 3A–C). This latter method can be used to enhace the speed of processing without a sacrifice in accuracy.

**Figure 3.**
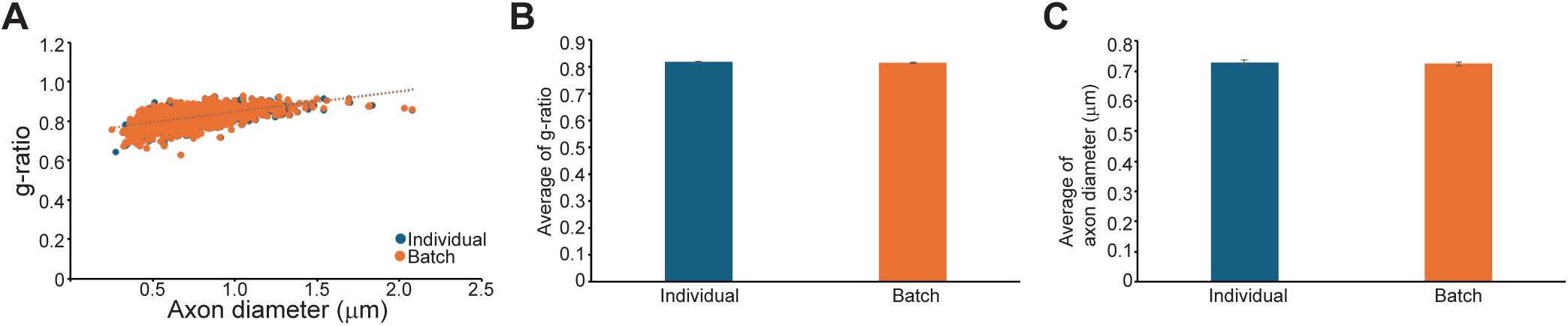
SnapG enables batch-wise g-ratio calculation equivalent to per-image analysis. A. Comparison of g-ratio values from per-image (Individual; n = 1027 axons) and batch-level (Batch; n = 1047 axons) calculations in P90 corpus callosum show both methods yield similar results. B. Bar graph showing average g-ratio calculated at individual image (0.819 ± 0.001) and batch-level (0.817 ± 0.001). C. Bar graph showing average axon diameter calculated at individual image (0.729 ± 0.007 μm) and batch-level (0.724 ± 0.007 μm).

## Discussion

G-ratios have been measured by different methods in multiple studies over the years. Variations in the imaging modalities used (e.g., optical vs. electron microscopy, MRI), in the fixation protocols employed, and the range of axon diameters selected for quantification, can all impact values obtained (Fields and Dutta, 2025; Gow, 2025). We developed SnapG to enable accurate and efficient g-ratio quantification from electron micrographs which are typically used to assess pathological changes in myelinated axons. SnapG incorporates a computationally efficient, unsupervised algorithm for segmenting myelinated axons and myelin sheaths, which we have integrated into an intuitive GUI application. The segmentation algorithm is entirely based on traditional image processing, offering speed and computational efficiency compared to deep-learning methods. The GUI application was also designed to process batches of images and streamline the workflow for manually removing false positives. Although SnapG does not include a built-in feature for manual segmentation of false negatives, it provides a set of tunable parameters that the user can adjust to minimize missed detection.

Among the practical advantages of SnapG are its streamlined batch processing capability and broad image format compatibility, resulting in a substantial reduction in analysis time. SnapG outputs were also highly consistent with g-ratios determined by manual tracing. The precise matching of SnapG to manually measured g-ratios is aided by choosing an appropriate percentile. As noted, SnapG traces the inner contour of each axon and, at every point along this contour, identifies the nearest white pixel located outside it. The user-selected percentile of these distances is then used as the representative myelin thickness. In our experience, the 60^th^ percentile optimized the correspondence between SnapG and manual measurements while minimizing errors in the PNS. The specific percentile to be used in an individual study can be determined empirically if desired by comparison to a small set of manually measured profiles that serve as controls.

Despite these advantages, SnapG does have limitations that stem from its reliance on threshold-based detection and contour segmentation, features common to many g-ratio algorithms. While inner contours are indicative of axonal boundaries in PNS (Fig. 1A), the CNS occasionally exhibits a periaxonal space-containing adaxonal myelin-between the axon and its surrounding myelin sheath (Fig. 2B, C). This structural feature complicates automated g-ratio quantification using SnapG. In such cases, users must either exclude axons with periaxonal space from analysis or manually measure them using custom contour definitions if inclusion is desired. In contrast to SnapG, MyelTracer is capable of measuring inner, outer, and axonal diameters, enabling the software to separate the periaxonal space (Kaiser et al., 2021). However, as SnapG does not detect axonal contours automatically, axons must be manually traced when a significant periaxonal space is present. In this study, 24.61 ± 2.16% of myelinated axons in P90 optic nerves required manual tracing. While this portion may differ depending on region and age, the full automation of all other steps does enable SnapG to significantly reduce the time required for g-ratio analysis.

A series of images captured under consistent brightness and contrast settings can be quantified simultaneously. However, a single threshold must be applied across all images, which may introduce variability. We evaluated both single-batch processing and multi-batch processing, and users may choose either approach depending on the consistency of their imaging conditions. Batch-processing markedly improves the efficiency of g-ratio analysis across multiple animals. Given SnapG’s reliance on threshold-based detection, some visually identifiable myelinated axons may fall below the threshold. To obtain accurate counts of myelinated axons per FOV, these undetected axons must be manually added.

In summary, SnapG represents a new method for g-ratio quantification that significantly reduces analysis time with overall strong correlation to manual quantification. SnapG provides a practical and efficient tool for g-ratio quantification, particularly in large-scale studies where high-throughput analysis is required; it is the method of choice in our own lab. One limitation shared by this and similar programs that measure g-ratios, is the inability to reliably detect and quantify unmyelinated axons and thereby quantify the percentages of myelinated vs unmyelinated axons. This is the result of the relatively low contrast of unmyelinated axons in many EM images. As AI technology continues to advance rapidly, we anticipate that tools capable of detecting unmyelinated axons will become available and serve as an adjunct to SnapG measurements.

---

**Supplemental Figure 1. Image processing workflow in SnapG**

A. Raw input image. B. Generated contour mask. C. Distance transform of the mask with the contour overlaid for visualization. D. Exclusion mask used to remove irrelevant regions. E. Outline mask delineating the target structure. F. Visualization of the exclusion mask applied to the distance transform. G. Visualization showing the *n* pixels closest to the current contour. H. Comparison of manual g-ratio measurements with g-ratios derived from the 30th–70th percentiles.

## Acknowledgements

We used Microsoft Copilot to assist with editing of the original draft.

