## Supplementary figures and images for "SnapG: An Automated Tool for Myelin g-ratio Measurement with Batch Processing Capability"

### Supplemental Figure 1

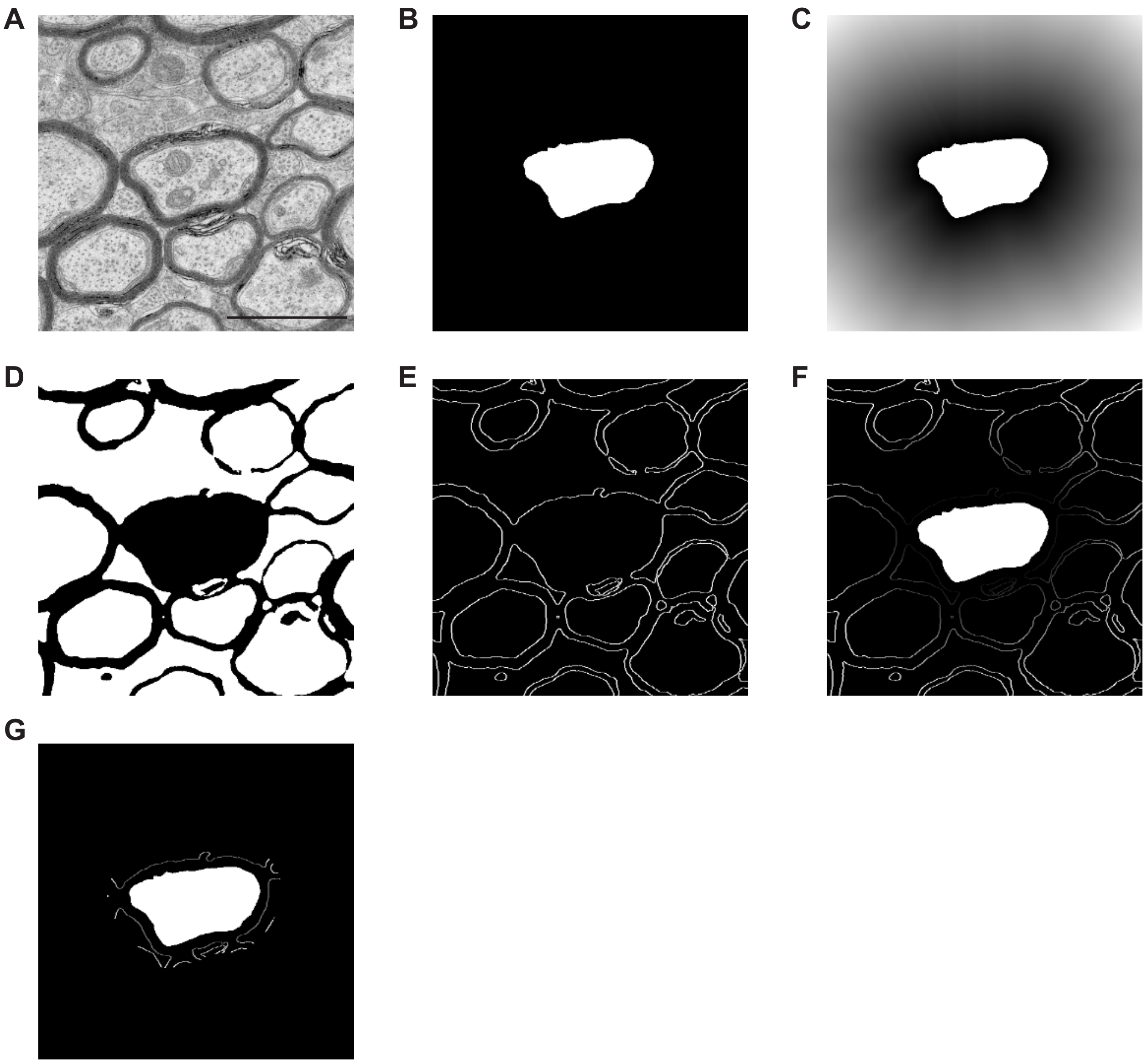


H

| | manual | 30 | 40 | 50 | 60 | 70 |
|---|---|---|---|---|---|---|
| mean + SEM | 0.669 + 0.004 | 0.698 + 0.003 | 0.692 + 0.003 | 0.682 + 0.003 | 0.675 + 0.003 | 0.667 + 0.003 |

Figure S1
